# Effects of nectar and sugar on the longevity of the mosquito *Anopheles gambiae* s.l.

**DOI:** 10.1101/2025.03.02.641044

**Authors:** Khadidiatou Cissé-Niambélé, Jacob C. Koella, Benjamin Koudou Guibéhi

## Abstract

In addition to taking blood-meals, mosquitoes regularly feed on the nectar of plants. The nectar, in particular the sugar it contains, serves as a source of energy that underlies many life-history traits, including longevity. To understand better how different nectars and sugars influence the longevity of mosquitoes, we allowed adult *Anopheles gambiae* s.l. to feed on several plant species (the flowers of *Thevetia nerifolia*, *Mandalium coromandelianum*, *Ixora coccinea*, and *Tabernanthe iboga*, and the fruit of *Carica papaya*), measured the concentrations of sucrose, fructose and glucose in the nectar and fruit juice, estimated the size of their sugar meal and measured their longevity. The plant that the mosquitoes fed on affected their longevity (which ranged from an average of 8.2 days when they fed on *C. papaya* to 21.1 days when they fed on *M. coromandelianum*), and the mosquitoes took larger meals (in a separate experiment) from the plants that gave a longer life. However, longevity was only slightly affected by the concentrations of glucose, fructose, sucrose, or total sugar in the sugar meals. In contrast, when we let mosquitoes feed on experimental sugar solutions consisting of glucose, sucrose, fructose, or trehalose at concentrations of 1.97 or 19.97 kcal per 100 ml, both the type and the concentration of sugar affected longevity. The mosquitoes lived approximately one week longer when fed sugar at the higher concentration and they lived longest (14.1 days) when fed with sucrose and shortest (4.8 days) when fed with trehalose. Overall, our results show the importance of nectar and sugar on the longevity of mosquitoes but suggest also that non-sugar components of nectar may have a large impact.

## Introduction

While female mosquitoes use blood meals for egg maturation, both sexes rely on various sources of sugar for survival after emergence [1] [2]. The availability and quality of sugar sources can strongly affect the vectorial capacity of mosquitoes [1] [3] [4]. One of the main sources of sugar for mosquitoes is the nectar of many plant species. Nectar is composed of different types of sugar (mainly sucrose, glucose, or fructose), amino acids, secondary metabolites, and microorganisms [5] [6]. Several studies have highlighted the impact of nectar on the life-history traits of mosquitoes, and have assumed that this impact is due to the sugars it contains [7] [8] [9] [10]. Since the type and concentration of sugars in nectar varies considerably among different plant species [6], this is expected to affect the life history traits of mosquitoes differently. However, there is currently limited information on how the size of the nectar meal and the types and concentration of sugars in nectar can affect life history traits of mosquitoes and, thus, their ability to transmit infectious diseases [3].

An important trait underlying the transmission of diseases like malaria is longevity, for infected mosquitoes can only transmit the malaria parasite, if they survive long enough for it to reach its infective stage. Longevity is influenced by adult feeding, as the energy available at emergence needs to be replenished by exploiting different sugar sources [1]. Additionally, the longevity of *Anopheles gambiae*, one of the main species transmitting malaria in sub-Saharan Africa, varies among the plant species it feeds on and it prefers to feed on the plant species that allow it to live the longest [8] [9] [10].

To increase our knowledge of this important aspect of mosquito ecology, we asked with two experiments whether the size of the nectar-meal and the concentrations of the sugars in the nectar influence the longevity of *A. gambiae* feeding on different plant species. First, we allowed mosquitoes to feed on the nectar or fruit juice of five different plant species that grow naturally in this locality and measured their longevity and the sizes of their nectar meals. We also measured the concentrations of sugars in the nectar or fruit juice of each plant. Second, we allowed mosquitoes to feed on experimental sugar solutions consisting of different types of sugars at two concentrations and assessed their longevity.

## Methodology

The experiments were carried out in an insectary in Agboville, Côte d’Ivoire, which was maintained at 25°C ± 2 °C, 76% ± 2 % relative humidity and a 12:12 h light: dark cycle. We used the offspring of *A. gambiae* that we had collected as larvae from irrigated rice fields in the town of Tiassalé, Côte d’Ivoire, where most mosquitoes are highly resistant to all four classes of insecticides [11]. The larvae were brought to the insectary and reared to adulthood. Their offspring were reared in trays containing between 120 and 150 larvae in 700-800 ml of tap water and fed daily with Tetramin baby fish food (TetraMin® tropical flakes, Tetra®, Blacksburg, VA, USA) according to their age [12].

### Plant-based diet

In a preliminary study, we tested whether mosquitoes fed on the flowers or fruit of 12 common species (Table 1), and selected for our experiments the five species on which mosquitoes were most likely to feed: the flowers of *Ixora coccinea*, *Malvastrum coromandelianum*, *Tabernanthe iboga*, and *Thevetia neriifolia*, and the fruit of papaya (*Carica papaya*).

**Table 1:** List of Plant Species used in the preliminary study.

| Plant species | Types of the plant species | Part of the plant used |
| --- | --- | --- |
| Tabernanthe iboga | Ornamental | flower |
| Thunbergia erecta | Ornamental | flower |
| Ixora coccinea | Ornamental | flower |
| Costus woodsonii | Ornamental | flower |
| Allamanda cathartica | Ornamental | flower |
| Thevetia nerifolia | Ornamental | flower |
| Lantana camara | perennial | flower |
| Mandalium coromandelianum | perennial | flower |
| Aspilia africana | perennial | flower |
| Centraterum intermedium | perennial | flowers |
| Carica papaya | seasonal | fruits |
| Mangifera indica | seasonal | fruits |

In the nectar or fruit juice of these five species we measured the total concentration of all sugars and the concentrations of glucose, fructose and sucrose. We extracted the nectar from the small flowers of *I. coccinea* and *T. iboga* using a syringe, and the nectar of *T. nerifolia* was extracted by pressing the flower into an Eppendorf tube. The fruit juice of *C. papaya* was extracted by crushing the pulp, allowing it to settle, and then collecting the juice from the top. We could not extract nectar of *M. coromandelianum* because we did not see any nectar in the flower, even on the flower stems, a small amount of sticky substance was occasionally visible, but could not be collected. In each sample, we measured the concentration of fructose using a fructose assay kit (Sigma Aldrich, Saint Louise, USA; product code FA-20), reading the optical density with a spectrophotometer (absorbance at 340 nm) (Thermo Scientific, Genesys 10S UV-VIS). We then determined the proportions of fructose, sucrose, and glucose in the samples with nuclear magnetic resonance spectroscopy (Bruker Avance Neo Ascend 600MHz, Bruker, Germany; Mnova NMR software package v.14.2.0, MestReLab Research S.L., Spain) [13] and calculated the concentrations of glucose, sucrose and total sugar from the concentration of fructose.

For NMR analysis, 50 µL of each nectar sample was dissolved in 450 µL of D₂O. Spectra were acquired using a Bruker Avance Neo Ascend 600 MHz spectrometer (Bruker, Germany), employing the noesygppr1D pulse program for water-suppressed ¹H NMR spectra. A total of 256 scans were recorded at 25 °C. Spectral processing was performed with Mnova software (version 14.2.0, MestReLab Research S.L., Spain). Sucrose, fructose, and glucose were identified in good agreement with published reference data [14] [15] [13].

Male and female mosquitoes were maintained separately in mesh-covered cages (30 × 30 × 30 cm). For each plant species we used nine cages for females and nine cages for males, each with 15 to 25 mosquitoes. Each cage contained one plant species as a sugar source, either as a number of flowers that would provide about 3 ml of nectar (placed in a 500 ml Erlenmeyer flask filled with tap water, plugged with cotton wool, and sealed with Parafilm) or a papaya fruit cut in half. Since for *M. coromandelianum* we could not extract and quantify any nectar, we provided 20 flowers of this species as a nectar source.

We used three of the cages per plant species and sex to assess longevity. From emergence onwards, the mosquitoes were provided access to the sugar source ad libitum, which was replaced daily. The cages were checked daily for dead mosquitoes until all had died.

We used the other cages to estimate the size of the sugar meal. For the first 48 hours after emergence, we provided the mosquitoes with water only and then placed them into cages with their sugar source and froze them at −20°C, 18 hours later. We estimated the size of the sugar meal from the relationship between wet weight (measured to a precision of 1 μg) and wing length (measured from the distal end of the alula to the tip with the image-processing software ImageJ). To do so, we first found the mosquitoes that had no sugar in their gut with an Anthrone test [16], and calculated the linear relationship between their wet weights and their wing lengths. We used this relationship to predict from their wing lengths the wet weight that a mosquito would have had, had it been unfed. The difference between the observed weight and the predicted weight (had it not fed) was used as a measure of the size of the sugar meal. We used the average size of the sugar meal taken from each plant species as a measure of their ability to feed on that species.

### Sugar-based diet

A sugar meal consisted of sucrose, fructose, glucose, or trehalose at a concentration of either 1.97 kcal or 19.7 kcal / 100 ml water and was provided on a wad of cotton placed on top of the cage.

Female mosquitoes were kept in three mesh-covered cages per diet (15 × 15 × 15 cm), each with 15 to 22 mosquitoes. The sugar meal was renewed daily, and mosquitoes were checked daily until they had all died.

### Statistical Analyses

The statistical analyses were performed using the software R version R-4.3.2. All analyses were linear mixed models, performed with the function lmer. We found the significance of the effects with the function **Anova** (package car), using a type 3 SS if the interactions were significant and a type 2 SS if they were not.

#### Plant-based diet

First, we assessed the impact of plant species on longevity with an analysis of variance that included plant species, sex and their interaction as fixed factors and cage as a random factor. We analysed the square-root of longevity so that the residuals were close to normally distributed. Second, we assessed whether the size of the sugar meal (i.e., the difference between the observed weight and the weight predicted for an unfed mosquito of the same size) differed among plant species with an ANOVA of the size of the sugar meal that included the same factors. Third, we assessed whether the size of the sugar meal affected longevity with an analysis of covariance (ANCOVA) of the square-root of longevity that included the average size of the sugar meal taken from a given plant species as a continuous variable, sex and their interaction, and cage as a random factor. We then assessed whether the sugar content of the plants affected longevity, ignoring data with *M. coromandelianum*, from which we were unable to extract nectar. We analysed the square-root of longevity first with an ANCOVA that included the total sugar content of the plants as a categorical variable, sex and their interaction, and then with an ANCOVA that included the content of each sugar (fructose, sucrose and glucose), sex, and the interactions between sex and each sugar. Both analyses included the cage as a random factor.

#### Sugar based-diet

We assessed the impact of individual sugar diluted in water on longevity of female mosquitoes by including types of sugar, concentrations and their interaction as fixed factors, and cage as a random factor.

## Results

### Plant-based diet

The concentrations of total sugar in the fruit juice or nectar of the plant species (excluding *M. coromandelianum*, from which we could not extract any nectar) ranged from 30.1 g/l for *C. papaya* fruit juice to 951.0 g/l for *T. iboga flower* nectar. In the three nectars, sucrose was the dominant sugar, but in the juice of *C. papaya* about a third of the sugar was fructose (**Table 1**).

**Table 2.** Concentrations (g/l) of sugars in the fruit of *C. papaya* and the three plants from which we could extract nectar.

| Plant species | Sucrose | Fructose | Glucose | Total sugar |
| --- | --- | --- | --- | --- |
| *T. iboga* | 882 | 41 | 22.0 | 951 |
| *I. coccinea* | 466.7 | 7.0 | 3.5 | 500.0 |
| *T. nerifolia* | 96.2 | 3.0 | 2.5 | 107.1 |
| *C. papaya* | 0.1 | 12.6 | 10.3 | 30.1 |

The average longevity of the 500 mosquitoes (242 males and 258 females) was 15.9 days and ranged from 8.2 days (7.21-9.33) when the mosquitoes had fed on the fruit of *C. papaya* to 21.1 days when they had fed on *M. coromandelianum* (19.1-23.3), (***χ***²=25.94, df=4, p<0.001) (**Fig. 1A**). Neither sex (***χ***²=0.42, df =1, p=0.516) nor the interaction between sex and plant species (***χ***²=2.55, df=4, p=0.635) affected longevity.

**Figure 1:**
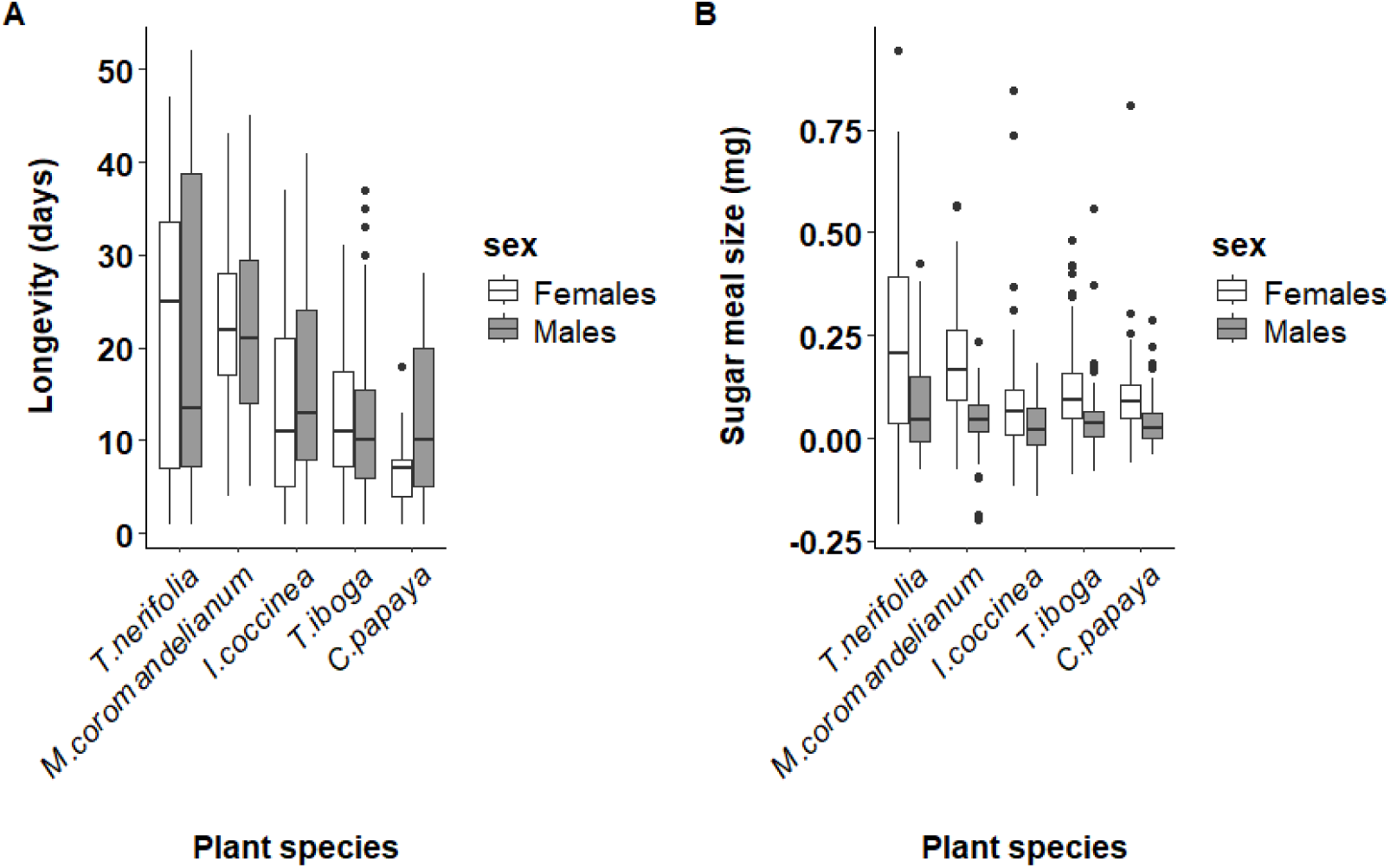
The effect of plant based diet on the longevity (A) and (B) the size of the sugar meal of mosquitoes. The rectangle represents the interquartile range, that is, longevities between the 25th and the 75th percentile. The thick horizontal line within the rectangle denotes the median value. The vertical lines span four times the interquartile range, and the dots show outliers that are beyond this range.

The average sugar meal size (so, the difference between the observed weight and the weight predicted for an unfed mosquito with the same wing length) ranged from 0.01 mg (−0.1-0.03 mg) for males that had fed on *I. coccinea* to 0.21 mg (0.18-0.25 mg) for females that had fed on *T. nerifolia*. Females took larger meals than males (ꭓ^2^=64.5, df=1, p<0.001), and the size of the sugar meals differed between the plant species (ꭓ^2^=41.29, df=4, p<0.001) (**Fig. 1B**). The effect of total sugar concentration has a slight effect on the longevity of mosquitoes (12=11.33, df=3, p=0.010), the lowest sugar level (30.07 g/L) was associated with the shortest mean longevity 8.24 days (7.2–9.3). Longevity increased markedly at 107.14 g/L, reaching its maximum 18.74 days (15.4–22.4). However, at much higher sugar concentrations (500 and 952.55 g/L), longevity decreased again to values 12.3 days (10.5–14.4); and 11.4 days (9.8–13.1), respectively comparable to those observed at the lowest concentration. But this effect was not influenced by sex (ꭓ^2^ = 0.63, df=3, p=0.427) (**Fig. 2**). Mosquitoes lived longer when they fed on plants from which they (in a separate experiment) took, on average, a larger sugar meal (ꭓ^2^=12.94, df=1, p<0.001), regardless of the sex of the mosquitoes (interaction sex*meal size: ꭓ^2^=0.01 df=1, p=0.916) (**Fig. 3**).

**Figure 2:**
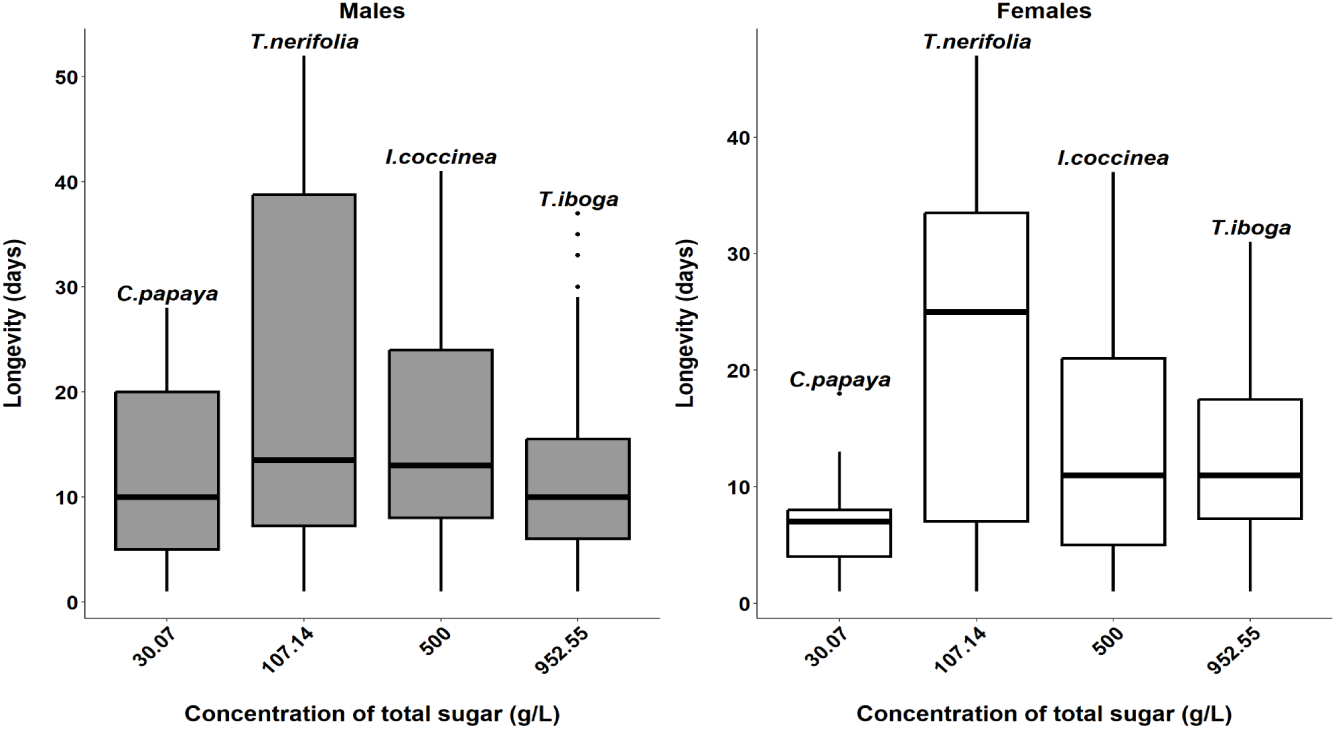
The effects of the concentration of total sugar in a plant sugar meal source on the longevity of mosquitoes. The rectangle represents the interquartile range, that is, longevities between the 25th and the 75th percentile. The thick horizontal line within the rectangle denotes the median value. The vertical lines span four times the interquartile range, and dots show outliers that are beyond this range.

**Figure 3:**
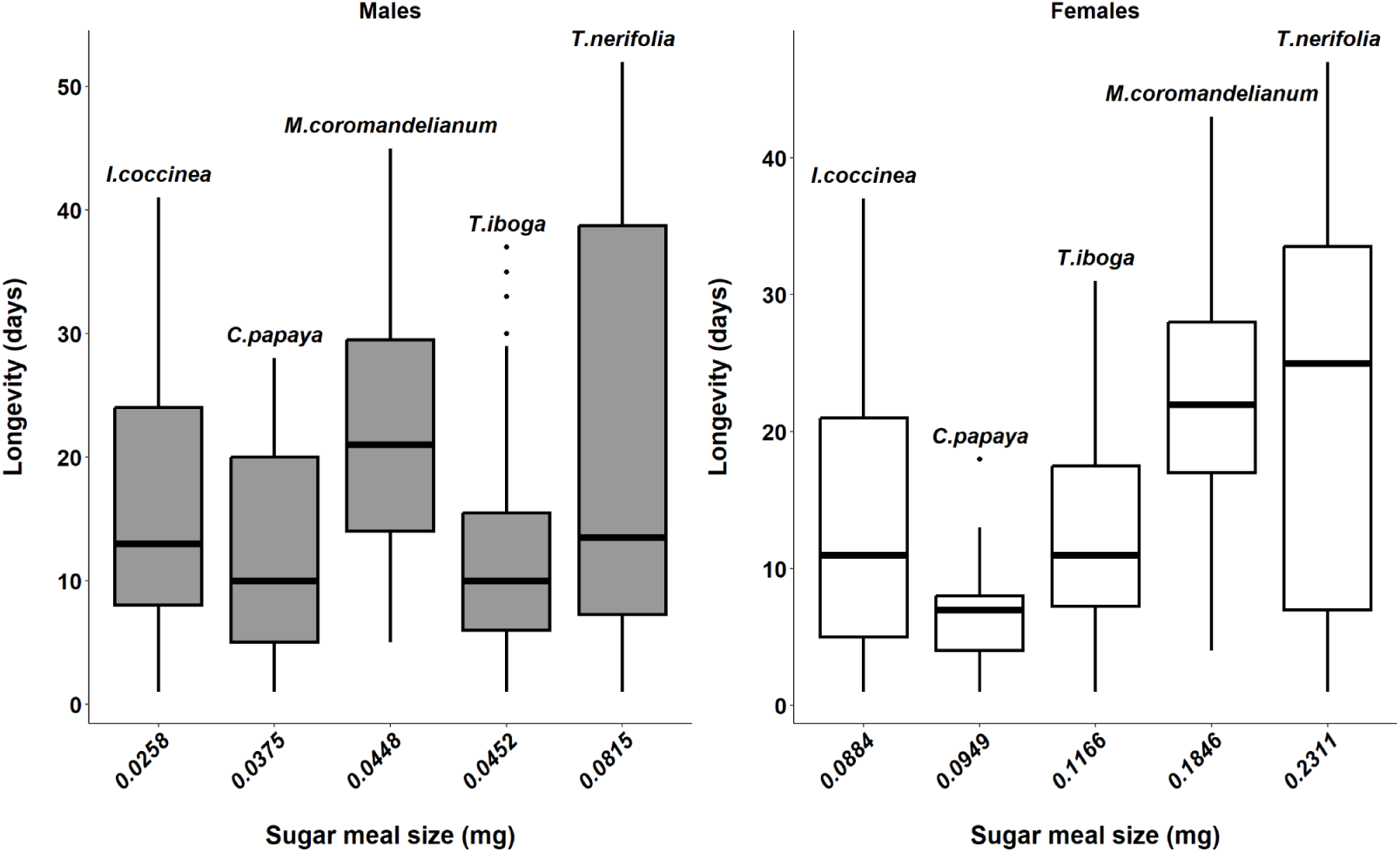
Relations between longevity and nectar meal size of mosquitoes feeding on different plant species. The rectangle represents the interquartile range, that is, longevities between the 25th and the 75th percentile. The thick horizontal line within the rectangle denotes the median value. The vertical lines span four times the interquartile range, and dots show outliers that are beyond this range.

### Sugar-based diet

The average longevity of the 406 female mosquitoes was 11.4 days. The type of sugar had a significant effect on the longevity of the mosquitoes (***χ***2=29.76, df=3, p<0.001); longevity was 14.1 days (12.7-15.6 days) if the mosquitoes fed on sucrose, 10.5 days (8.6-12.5) if they fed on glucose, 11.7 days (10.3-13.2) if they fed on fructose and 4.8 days (4.3-5.7) if they fed on trehalose. The higher concentration of sugar increased longevity from 6.6 days (5.8-7.4) to 13.4 days (12.2-14.7) (***χ***2=25.99m df=1, p<0.001).

The effect of sugar concentration on longevity varied greatly depending on the type of sugar ingested by the mosquitoes (***χ***2=19.10, df=3, p<0.001). For trehalose, an increase in concentration from 1.97 Kcal/100ml water to 19.7 Kcal/100 ml of water resulted in a marked increase in longevity from 2.3 (2.1-2.5) to 8.6 days (7.1-10.3) and for glucose it resulted in an increase from 4.1 (3.3-5.0) to 17.8 days (15.0-20.8). For fructose and sucrose, the effect of concentration was more moderate (for fructose, 11.3 (9.9-12.8) to 12.1 days (12.5-15.1); for sucrose, 12.8 (11.0-14.6) to 15.4 days (13.3-17.6), (**Fig. 4**).

**Figure 4:**
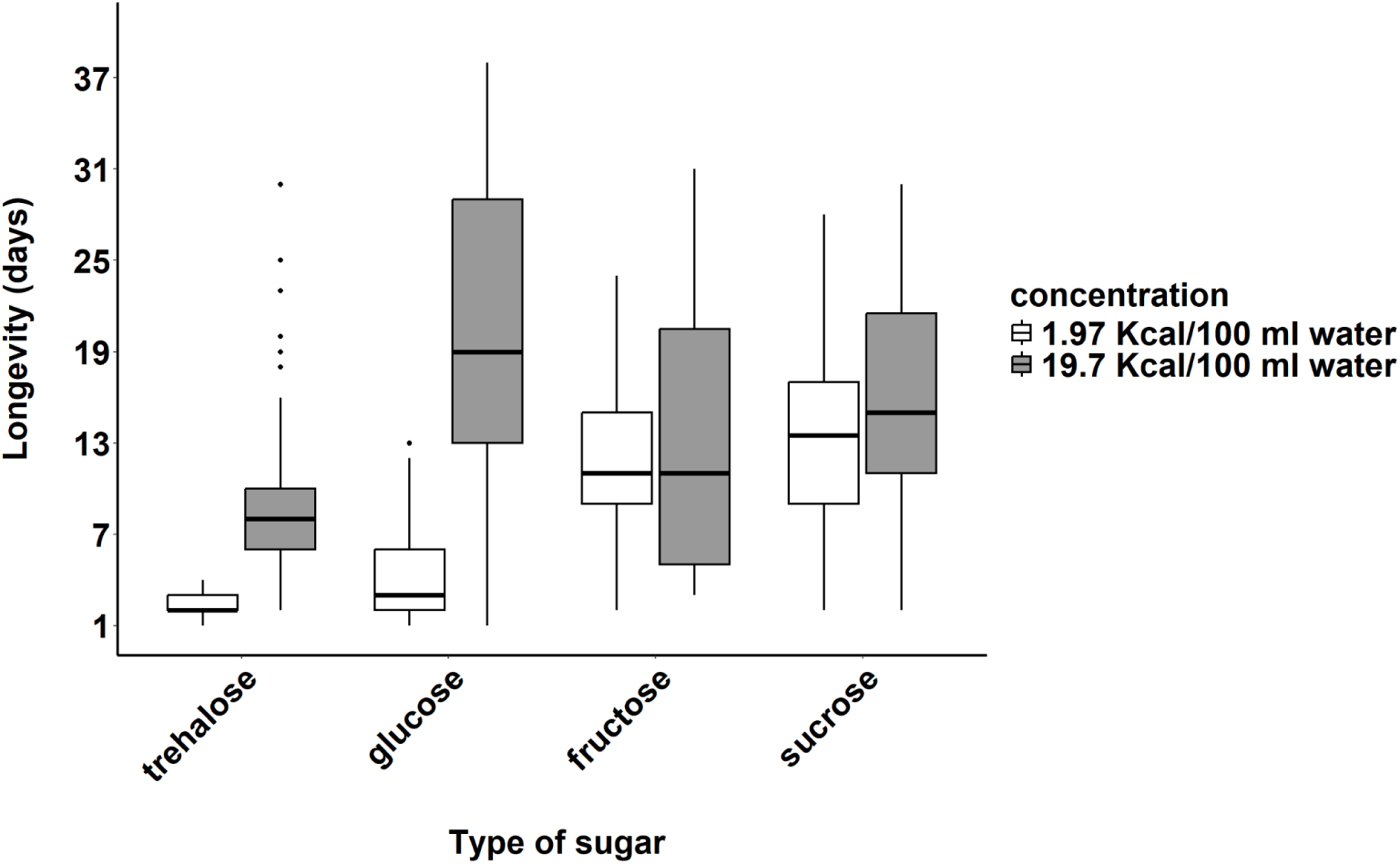
Effects of types and concentrations of sugar on the longevity of mosquitoes; The rectangle represents the interquartile range, that is, longevities between the 25th and the 75th percentile. The thick horizontal line within the rectangle denotes the median value. The vertical lines span four times the interquartile range, and the dots show outliers that are beyond this range.

## Discussion

In our experiments the type and concentration of sugar in sugar-meals and the species of plant providing nectar strongly affected the longevity of *A. gambiae* sl.. However, the concentration of sugars alone seems to exert a slight influence on longevity suggesting that other components in nectar strongly influence longevity.

The concentration and type of sugar that mosquitoes feed on have strong consequences for their life-history [17] [18]. In our experiment, mosquitoes fed with a high concentration of sugar lived seven days longer than those fed with a 10-time lower concentration, and mosquitoes lived longer when fed a sucrose meal than when fed the other sugars, in particular at the lower concentration. This corroborates the findings of a previous experiment in which mosquitoes fed with sucrose accumulated more energy reserves than those fed with glucose or fructose [19] in providing an energy supply and allowing stress adaptation in mosquito hemolymph [20] [21] [22]. It corroborates, also, a study by Cissé, Koudou and Koella (in prep.), in which mosquitoes fed with trehalose were more likely to die sooner after exposure to deltamethrin than those fed on other sugars. One reason may be that the metabolism of trehalose is not efficient if glucose is lacking, which leads to a deficiency in ATP [23] [24] [25].

The differences in longevity of the mosquitoes feeding on different plant species align with previous findings [8] [10,26] [27], in particular increased longevity when feeding on *T. nerifolia* [7] and the shorter life when feeding on *Carica papaya* [10,26].

The main component of nectar is sugar, but the differences we observed among plants was slightly linked to their sugar content. Indeed, neither the total sugar concentration nor the concentration of the individual sugars explained strongly the difference in the longevity. While this result contrasts with studies showing a positive correlation between sugar content in plant parts and mosquito survival [9] result was found in *Conopomorpha sinensis*, where additional honey in the diet had no significant effect on longevity [28]. This suggests that other components than sugar, for example amino acids and secondary metabolites, play an important role in the longevity of mosquitoes [29]. Indeed, the amino acids contained in nectar have been shown to increase the longevity of female mosquitoes *Culex quinquefasciatus* [30].

The variation in nectar intake among plant species suggests that mosquitoes may prefer some food sources over others. Such preferences can be linked to physical traits like the brightness of flowers or the shape of the corolla or the position of nectaries, which would affect the ability of mosquitoes to access the nectar [31]. They can also be linked to chemical emissions of the plants and nectars, which in turn can be linked to the quality of the nectar [9]. Indeed, plants that are preferred by mosquitoes provide more energy reserves than the less preferred [32]. This in turn leads to a positive correlation between nectar intake and longevity, for mosquitoes feeding on more nutritious plants benefit from increased sugar for energy and vital functions and therefore live longer [33] [34].

## Conclusion

The longevity of mosquitoes varies depending on the plant species from which they obtain a sugar meal although the concentration of sugar alone in the nectar does not strongly explain this longevity. However, different concentrations of sugar diluted in water have a significant impact on longevity. This result highlights the fact that there may be other compounds in the nectar that have a major influence on the longevity of the mosquitoes observed.

## Acknowledgements

This work was supported by SNF grant 310030_192786 and the donation fund of the University of Neuchâtel. We thank Professeur Koné Mamidou and his technician Mrs Dougouné bi Goré for their assistance for the identification of plant species. We would like to thank all the members of the Laboratory of Ecology and Epidemiology of Parasites at the University of Neuchâtel. The chemistry laboratory of the University of Neuchâtel, particularly Dr. Sylvain Surtour and Dr. Gaussan Gaëtan, for their assistance in determining the sugar proportions in nectar using NMR. We would like to thank Professeur Ouattara Foungoye Alassane for their advances in scientific research in general and in statistical analyses. We would like to thank Mrs Kouamé Romuald for his assistance during field work. We also thank the families who allowed us to collect their flowers and the technicians of the medical entomology laboratory at CSRS for their help.

